# A neural network-based framework to understand the Type 2 Diabetes (T2D)-related alteration of the human gut microbiome

**DOI:** 10.1101/2020.09.06.284885

**Authors:** Shun Guo, Haoran Zhang, Yunmeng Chu, Qingshan Jiang, Yingfei Ma

## Abstract

To identify the microbial markers from the complex human gut microbiome for delineating the disease-related microbial alteration is of great interest. Here, we develop a framework combining neural network (NN) and random forest (RF), resulting in 40 marker species and 90 marker genes identified from the metagenomic dataset D1 (185 healthy and 183 type 2 diabetes (T2D) samples), respectively. Using these markers, the NN model obtains higher accuracy in classifying the T2D-related samples than machine learning-based approaches. The NN-based regression analysis determines the fasting blood glucose (FBG) is the most significant association factor (P<<0.05) in the T2D-related alteration of the gut microbiome. Twenty-four marker species that vary little across the case and control samples and are often neglected by the statistic-based methods greatly shift in different stages of the T2D development, implying that the cumulative effect of the markers rather than individuals likely drives the alteration of the gut microbiome.

## BACKGROUND

The gut microbiota, which inhabits the human intestinal tract, is a complex ecosystem consisting of 10^14^ microbial cells (Tropini, Earle et al. 2017) and it has been identified playing a critical role in a variety of human diseases (Morgan 2012, LM and MJ 2015, Manrique, Bolduc et al. 2016). Metagenomic studies have provided great opportunities to get valuable insights into how the gut microbiota is associated with various human diseases via various well-developed bioinformatic tools and algorithms.

In the field of microbiome research, a common practice is to utilize the statistics-based strategies (e.g., Spearman’s correlation coefficient, Wilcoxon rank-sum test, etc.) to identify the biomarkers associated with the gut microbiome according to the disease states (Segata, Izard et al. 2011, Qin, Li et al. 2012, Karlsson, Tremaroli et al. 2013, Nielsen, Almeida et al. 2014, Liu, Hong et al. 2017, Wen, Zheng et al. 2017), for example, the software LEfSe (Segata, Izard et al. 2011, LA, CF et al. 2014, Yu, Guo et al. 2017). These methods can identify metagenomic features that have statistically significant differences between case and control groups. Nevertheless, these statistics-based analyses are typically based on the independent or linear assumption, whereas dysbiosis of intestinal flora is complex and likely depends on the nonlinear effects of many microbes (Ilseung and Blaser 2012). To this end, these methods may neglect some potential metagenomic biomarkers that might contribute to the disease-related alteration of the human gut microbiome but have no detectable statistic changes across the samples. For example, *Veillonella parvula* has been reported to be related to T2D (Koh, De et al. 2016, Zhao, Zhang et al. 2018), however, the abundance of which varied little across the samples (Qin, Li et al. 2012).

Machine learning has recently attracted growing attention in biological research. Some well-known algorithms including Support Vector Machine (SVM), Random Forest (RF), Hidden Markov Model (HMM), Bayesian network (BN) as well as Gaussian network (GN), have been used in prediction of the protein binding (Alipanahi, Delong et al. 2015), the metabolic functions in microbial communities (Sung, Kim et al. 2017), characterization of the transcriptional networks (Ghanat, Ung et al. 2017), human microbiome studies (M. Yazdani 2016, Pasolli, Truong et al. 2016, Cammarota, Ianiro et al. 2020), *etc*.. As machine learning can generate models and find predictive patterns from large datasets, it would impact microbiome research and other biology fields (Camacho, Collins et al. 2018). However, these machine learning algorithms have some limitations. For example, SVM is a linear model and HMM and BN depend on some probability-based hypotheses. To this end, some T2D related species, such as *Coprobacillus catus* (Koh, De et al. 2016, Zhao, Zhang et al. 2018) would not be identified by these methods since the abundance of the species varies little across samples (may not have the linear correlation with the disease) and the average abundance is very low (would be the outlier under some probability distribution) (Qin, Li et al. 2012).

Deep learning-based methods, which typically use the neural network (NN) with multiple hidden layers, show promising performance in recent studies as they can identify novel patterns that would have been ignored by other methods in a given complex dataset (Angermueller, Pärnamaa et al. 2016, Morton, Aksenov et al. 2019, Galkin, Mamoshina et al. 2020). Recently, these methods have been applied in the field of biology, such as predicting special genes/protein functions(Kulmanov, Khan et al. 2017, Arango-Argoty, Garner et al. 2018), identifying medical diagnoses(Kermany, Goldbaum et al. 2018), drug discovery(Chen, Engkvist et al. 2018), etc. However, so far, very few microbiome-related studies have utilized the technique. One reason may be that the number of available samples is limited in microbiome studies. The deep learning model typically having many layers and neurons requires massive samples for training. Another reason is that applying the NN model to the microbiome data is still a challenge due to its ‘black box’ nature. This makes it difficult to demonstrate which input feature plays a decisive role in the output.

Here, we propose a framework combining NN and RF algorithms for identifying the biomarkers in linking the gut microbiome with T2D based on the microbial profiles. To demonstrate the utility of our approach, we took advantage of two publicly available independent metagenomic datasets from Chinese diabetes patients and non-diabetic controls (Qin, Li et al. 2012). We first trained a NN model for predicting the disease state of the samples. Further, RF was used to rank the microbial features, and the most important features contributing to the prediction accuracy of the NN model were selected as microbial markers. Using these microbial markers, we finally constructed interaction networks and performed the regression analysis to understand the alteration of the human gut microbiome in the development of T2D.

## METHODS

### Datasets

The whole metagenomic sequencing datasets used in this study were generated in two studies, respectively (Qin, Li et al. 2012, Zhao, Zhang et al. 2018). The first dataset D1 (Qin, Li et al. 2012) (accession number in GenBank: PRJNA422434) is generated by shotgun sequencing of the DNA extracted from the stool samples from Chinese individuals (183 samples from the subjects with T2D and 185 control samples from the healthy subjects). The second dataset D2 (Zhao, Zhang et al. 2018) (accession numbers: PRJEB15179 and PRJEB14155) is obtained from the metagenomic shotgun sequencing of 391 samples from the Chinese patients with T2D at different days (0, 28, 56 and 84), where all samples are divided into two groups with two diets for studying the effects of the dietary on T2D. In this study, the dataset D1was used for training the Neural Network model and selecting the microbial markers. Since the dataset D2 has only positive samples (i.e., case samples), 30% control samples (n=56) of D1 were randomly selected as the control for D2. The remaining samples of D1 were assigned as dataset D1−. D2 and the added control samples (i.e., D2+) can be viewed as the dataset that is independent of D1−. The dataset D3 from the European women (Karlsson, Tremaroli et al. 2013) (accession number: ERP002469, 53 T2D samples, 43 control samples) was also used for testing. In the testing analysis, D1−was used for training the model, D2+ and D3 for testing.

### Species abundance and gene abundance calculation

The microbial profile of each sample on taxonomic species-level was estimated by the software MetaPhlAn2.7.4 (Segata, Waldron et al. 2012) based on the metagenomic sequencing read data and the relative abundance of each species was calculated. The species present in at least two samples of D1 were selected for profiling the D1, D2, and D3. The bacterial gene profile of each sample was estimated using the software Diamond (v0.9.14.115) against the COG (Clusters of Orthologous Groups) database with cut off e-value 1e-10 (Buchfink, Xie et al. 2015). The genes present in at least two samples of D1 were calculated for each sample for profiling D1, D2, and D3 on gene-level. The relative abundance of each gene was calculated based on the number of the metagenomic sequencing reads matching to the gene and subsequent normalization by the total read number of the sample.

### Neural network (NN) model training process and comparison

We used D1 samples to train the NN model and determined the parameter setting for the model based on the microbial profiles. There are no standard approaches to determine the parameters (e.g., the number of layers and the number of nodes in each layer) for a neural network. A common practice is to set up a range of parameters and choose the suitable ones, with which the NN model has the best performance. The datasets used in this study contain only hundreds of samples. Therefore, the number of hidden layers was set up in a range from 2 to 5, and the range of the number of nodes in a layer was set up from 5 to 25. The prediction performance (i.e., accuracy) of the NN model with different parameter values was assessed using 5-fold cross-validation (CV) based on the microbial profiles (species-level and gene level, respectively) of D1. In 5-fold CV, all samples of D1 were randomly divided into five equal size subsamples, and four were used for training, the remaining one for testing. The process was repeated five times, and each of the five subsamples was used once as the testing samples. To assess the performance of the model, we adopted 20 runs of 5-fold CV, resulting in a total of 100 trials for testing. We then chose the parameter settings with the best performance in this study. Rectified linear units (ReLU) were used for the hidden layers, and the softmax *function* was used in the output layer when we used the NN model for a classification task. Meanwhile, The loss function (cross entrop loss) was used for training. The code was implemented using Keras package in the Python (3.6). Briefly, let *S* = [*s*_1_, *s*_2_, …, *s*_*n*_] to be the abundance of the *n*th-marker in one sample, and prediction of its state is as follows:

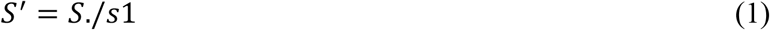

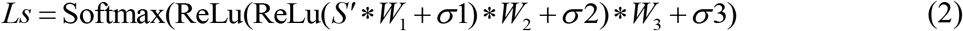

Where *s*1, *W*_1_, *W*_2_, *W*_3_, *σ*1, *σ*2, *σ*3 are the model parameters (Table S1 and Table S2), and Ls represents the predicted result of the sample (0 for the non-diabetic, 1 for the diabetic).

Meanwhile, the performances of four widely used classifiers including SVM-linear, RF, SVM-RBF (We used the built-in function *fitcsvm* of MATLAB software with setting ‘KernelFunction’ value to ‘linear’ and ‘rbf’ respectively), and KNN (the built-in function *fitcknn* of MATLAB software was used with setting the number of neighbors to five) were also evaluated in the same way for comparison. All these classifiers were implemented on the microbial profile of D1 on species-level. Their prediction performance metrics were averaged by 20 runs of 5-fold CV in the experiment.

### T2D-related marker species identification and test

We used RF and the NN model to identify the marker species associated with T2D. Briefly, the package of ‘RF_Class_C’ of the MATLAB software was used with default parameters on the species-level profile of D1 samples to calculate the scores (i.e., feature weights) of all the species according to their impacts on the classification output as random noise was added on them. Based on the scores, we ranked all the species of the samples in D1. Then, the feature selection process was implemented using the NN model on the ranked species-level profile of D1. Using 20 runs of 5-fold CV, the average accuracy of the trained NN model to classify the T2D-related samples using the profile of the ranked top-*k* (*k* ranged from 5 to 60) species of D1 was calculated. The minimum number *n* of the species with the highest average accuracy was determined and thus top *n* species were selected as T2D-related marker species contributing to the alteration of the human gut microbiome. Also, the prediction performance of the NN model (retrained on D1−dataset) with the selected marker species in classifying T2D-related samples was further assessed using the D2+ dataset. As a comparison, the marker species selected by the software LEfSe (Segata, Izard et al. 2011) were fed to the NN model (trained on D1−dataset) and the prediction performance was assessed using the D2+ dataset as well. Besides, we also tested the prediction performance of the NN model with the marker species (selected and trained on the D1 dataset) on the independent dataset of D3, which comes from European women.

### Marker species interaction network inference and analysis

To further investigate the relationship between the selected marker species, GENIE3 was utilized for constructing the species-species interaction network of the marker species. The package of ‘GENIE3_MATLAB’ of the MATLAB software was performed with default parameters to calculate the scores of the links between the marker species according to the correlation strength, and then the links were ranked based on the scores. The higher the score is, the more reliable the link between the marker species is. We also used a refinement procedure (Guo, Jiang et al. 2016) to improve the constructed network under the assumption that the links of the hub nodes would be more important and reliable. Once GENIE3 was implemented, an adjacency matrix *M* will be obtained, where *M*_*ij*_ represents the reliability level of the link from node *i* to node *j*. And the refined adjacency matrix 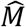 is given as:

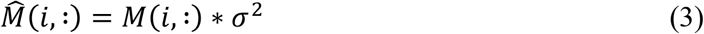

where 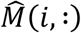 is the *i*-th row of *M* and *σ*^2^ is a variance in the *i*-th row of *M*. It should be noted that if many links come from the same node, the variance in a row of *M* corresponding to the node would be elevated. Finally, we selected the top 100 links in our experiments to reconstruct the interaction network.

### Marker gene identification and test

In the same way of species marker identification, we identified the marker genes by using RF and the NN model based on the gene profile of the D1 dataset. Briefly, the average prediction accuracy of the NN model using the profile of the top *k* (*k* from 5 to 100) genes of D1 was calculated using 20 runs of 5-fold CV. Then, we selected the top *n* T2D-related marker genes, with which the NN model has the highest average accuracy in classifying T2D-related samples. The performance of the NN model using the selected marker genes (trained in D1−dataset) in classifying T2D-related samples was further tested using the D2+ dataset. The functions of the marker genes were annotated by mapping the genes to the KEGG database. Also, the performance of the NN model with the marker genes (trained on D1 dataset) was tested on the dataset from European women (i.e., D3 dataset).

### Marker gene interaction network inference and analysis

Using GENIE3 and the above procedure of the marker species, the gene-gene interaction network was constructed based on the maker gene profile of the D1 dataset. Similarly, we selected the top 100 links of genes to reconstruct the interaction network.

### NN-based regression analysis of the human gut microbiome and the T2D-related factors

To investigate the correlations of the T2D-related factors (including fasting blood glucose (FBG), age, body mass index (BMI), and body weight) with the T2D-related alterations of the gut microbiota, the NN-based regression analysis was performed. The regression model for each factor was built using the same structure of our NN-based classification model (i.e., the number of hidden layers and the number of nodes in these layers) except for the output layer (one node for regression, without using activation function), and training based on the mean-square error. The values of each factor in D1 (i.e., the dependent variable) were predicted by the corresponding trained model with our marker species (the dependent variables) using 5-fold CV. Moreover, the correlation coefficient (i.e., Pearson’s linear correlation coefficient), as well as the corresponding P-value between the predicted values and the real values for each factor in D1 were calculated, which reflects the degree that the factor is influenced by the abundances of the marker species.

To explore how the markers altered along with the dynamic change of FBG in the development of T2D, we stratified all samples of D1 into four intervals according to the quantile statistics of the corresponding FBG values (i.e., Q1:< 5.02; Q2: 5.02∼6.21; Q3: 6.21∼8.8; Q4: > 8.8). Then, the average abundances of the marker species of the samples within the corresponding interval were calculated and normalized (mapped to (0, 1)).

### Accuracy, precision, recall, and F1 metrics

The three metrics were defined as follows:

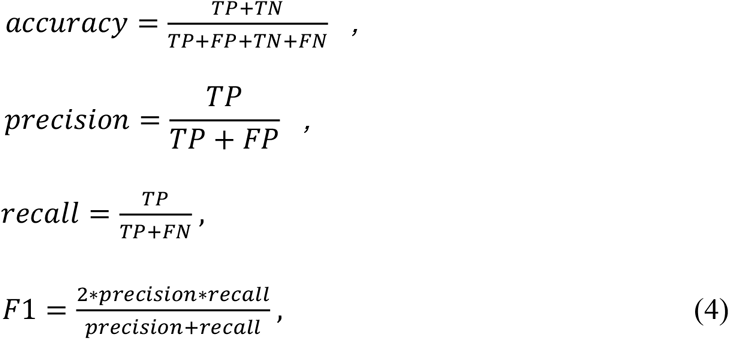

where TP indicates true positives (i.e., a case sample is predicted correctly), FP false positives (a control sample is predicted as a case sample), and FN false negatives (a case sample is predicted as a control sample). The accuracy measure reflects the overall quality of classification for the classifier. The precision measure focuses on how accurate the diabetic status predicted by the classifier is. The recall measure answers the question that what proportion of the diabetes samples could be identified by the classifier. And F1 measure is the integration of the precision measure and the recall measure.

### ROC (receiver operating characteristic) curve and AUC (area under the curve) metric

The ROC curve indicates the true positive rate (i.e., the number of correctly predicted case samples divided by the total number of samples predicted as case samples) against the false positive rate (i.e., the number of samples that wrongly predicted as case samples divided by the total number of samples predicted as control samples). The AUC metric ranges from 0.5 to 1 (the greater, the better), is computed from the ROC curve, which summarizes false positive and true positive rates. The AUC measure summarizes false positive and true positive rates and is robust to the unbalance situation of each outcome.

## RESULTS

All metagenomic samples were profiled on taxonomic species-level and functional gene-level, respectively. We first calculated the relative abundances of the species for each sample of D1, resulting in 270 species for analysis. The abundances of these species were calculated for all samples of D2 and D3. The genes encoded by the metagenomes were assigned using the COG database and the relative abundances of 4632 genes present in at least two samples were calculated for each sample for profiling D1 on gene-level. The relative abundances of these genes were calculated in the D2 and D3 datasets as well. All these data are available on the website (https://github.com/gsgowell/microbial_markers_identification).

### The NN model has higher performance in classifying T2D samples than other approaches

To train the NN model using the vector of the relative abundances of the gut microbial species of D1 samples in classifying human gut samples according to their T2D state, we first determined the parameters of the NN model. As shown in Figure 1a and Table S3, we find that the average accuracy of the NN model reaches 97.4% as the number of hidden layers is 2 and the performance is not improved with the increase in the number of hidden layers (Figure 1a). We also examined the effect of the number of nodes in each hidden layer (Figure 1a), and the result indicates that the best-average accuracy of NN can be achieved (98.4%) when the parameters are 18 and 9, respectively. Thus, we chose the parameter settings of 2 dense hidden layers of 18 and 9 nodes, respectively, for the NN model.

**Figure 1.**
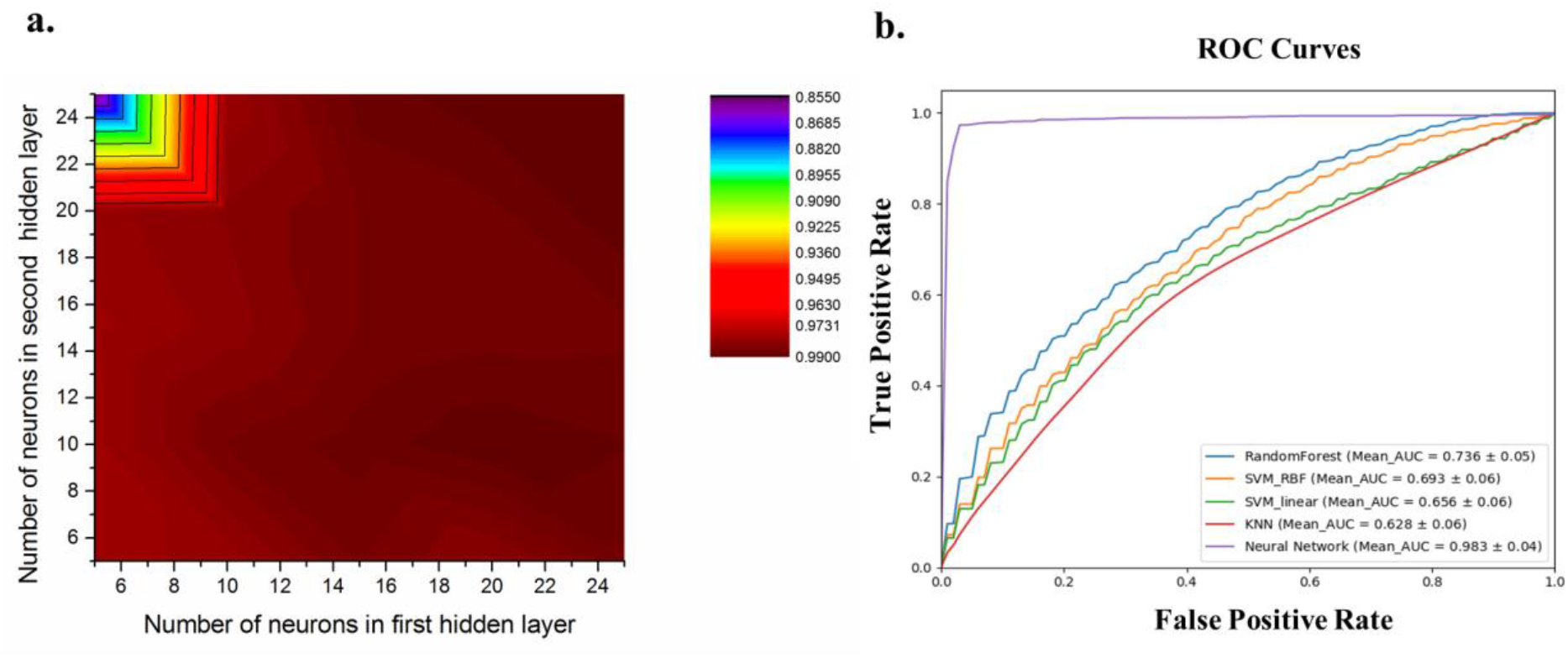
Neural Network performance evaluation and parameter selection. **a**. Average accuracy with different neurons based on the species level of D1. The results were obtained using 20 runs of 5-fold CV. The best performance was achieved as the parameters were set to 18 and 9 for the first and second hidden layers respectively. The colors show the prediction performances (i.e., accuracy) of the NN model with different parameter values. **b**. Average ROC curve of four classifiers (Neural Network, SVM-linear, SVM-RBF, KNN). The results were obtained by using 20 runs of 5-fold CV.

We assessed the performances of the NN model as well as SVM, SVM-RBF, RF, and KNN methods in classifying the T2D-related samples of the D1 dataset based on the microbial profile using 20 runs of 5-fold CV. The obtained results of the five methods with all metrics were listed in Table 1. The ROC curves were plotted in Figure 1b and thus the corresponding AUC metrics were calculated for each method. Compared with the other four methods, the NN model performes significantly better for all metrics (all are more than 98%), indicating the powerful nonlinear fitting ability of the NN model that likely can extract the most predictive features. The other four Methods have very similar results. For accuracy, precision, and AUC, all four classifiers (KNN, SVM-linear, RF, and SVM-RBF) could reach around ∼60%, similar to the results described in (Pasolli, Truong et al. 2016), much less than those of the NN model (Table 1). Also, they have lower performances on both recall and F1 (especially for KNN) than the NN model, which means that the proportion of the T2D samples that can be identified by these machine learning classifiers is low.

**Table 1.** Prediction performance comparison of four classifiers (mean ± standard deviation) on the T2D dataset.

| Classifier | Accuracy | Precision | Recall | F1 score | AUC |
| --- | --- | --- | --- | --- | --- |
| SVM-linear | 0.629±0.05 | 0.617±0.07 | 0.571±0.08 | 0.591±0.07 | 0.656±0.06 |
| SVM-RBF | 0.644±0.05 | 0.631±0.07 | 0.528±0.10 | 0.572±0.08 | 0.693±0.06 |
| KNN | 0.603±0.05 | 0.614±0.09 | 0.327±0.09 | 0.422±0.09 | 0.628±0.06 |
| Random Forest | 0.660±0.06 | 0.670±0.06 | 0.631±0.09 | 0.647±0.06 | 0.736±0.05 |
| Neural Network | **0.984±0.04** | **0.984±0.04** | **0.983±0.05** | **0.984±0.04** | **0.983±0.04** |

The abundance of the 270 species is used as the feature for the classifiers. Prediction performance metrics were calculated by 20 runs of 5-fold CV for all classifiers. In bold we report the best value for each prediction performance metric.

### The identified marker species play a decisive role in the T2D-related alteration of the human gut microbiota

We further investigated which species rather than all of the gut microbiota, play a decisive role in classifying the T2D-related samples using the NN model. We used the feature selection method-RF (Breiman 2001) for ranking all species of D1 samples according to their importance scores. We then used the NN model on top *k* (*k*=5,10,15,20,…,60) species with 20 runs of 5-fold CV to classify the samples. The prediction performances of the NN model using top *k* species were assessed according to their average accuracies in classifying the T2D-related samples. It can be seen from Figure 2a that when the top 40 species were selected for the NN model, the average accuracy reaches the highest value (**98**.**6%**). This result is almost the same as that of the model on all species (n=270) (Figure 2a and Table 1), suggesting that these species can be used to delineate the T2D-related alteration of the gut microbiota. Thus, we took the selected top 40 species as the marker species (Table S4).

**Figure 2.**
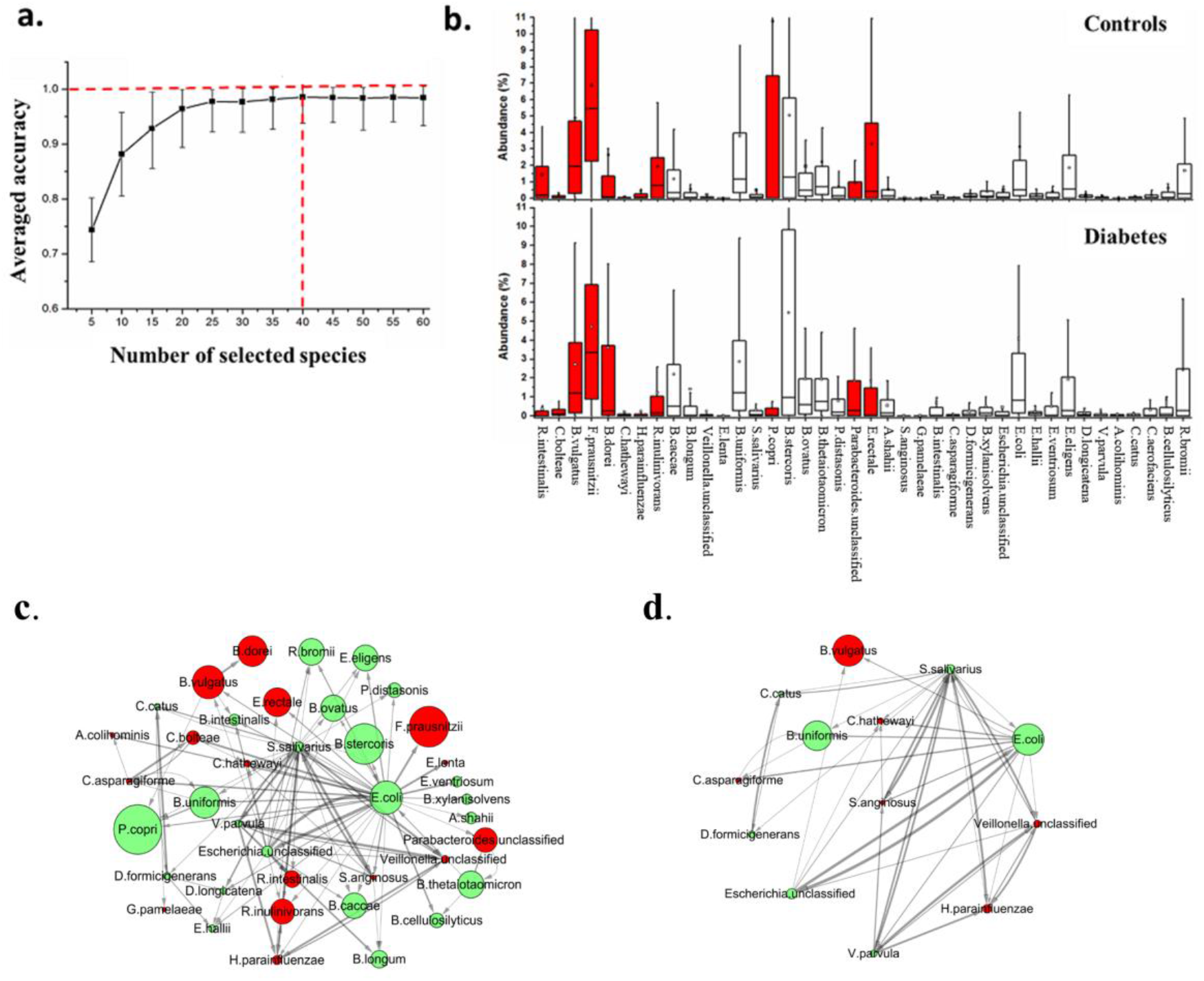
Marker species identification and interaction network construction. **a**. Average accuracy of classification with a different number of the ranked species based on Neural Network-based classifier by using 20 runs of 5-fold CV on the species-level profile of D1. And the peak value could be obtained as the 40 species were selected, thus these species were identified as the marker species in this study. **b**. The distribution of the abundance of marker species in the diabetes-related dysbiosis. A comparison between the cases (n = 183) and controls (n = 185) from the species-level profile of D1. Using the Kruskal-Wallis test, the species with red color were identified that the abundances varied significantly (P-value < 0.05) across all samples. The order of the marker species is the same as that in Table S4. **c**. The interaction network of marker species determined by GENIE3, where the top 100 most reliable interactions between marker species were selected. In the network, each node represents the respective marker species; the constructed internetwork is directed and the arrow between Nodes indicates the direction of the interaction, namely, each arrow from Node A to B means that the abundance of B is mainly affected by that of A. The width of the edges between nodes is proportional to the reliability of the linkage between the nodes. The size of each node is proportional to the average abundance of the marker species. The network layout was visualized by the **Cytoscape** software using a circular layout. The nodes with red color are the marker species that significantly are enriched in the diabetic samples or the control samples. **d**. Interactions between marker species with a degree of more than 5.

We further employed the independent dataset D2+ to assess the prediction capability of the NN model with the 40 marker species. In this test analysis, with these marker species, the species-level profile of D1−was utilized for retraining the NN model, while D2+ was used for testing. The accuracy in classifying T2D samples of D2+ reaches **85**.**2%**. We also chose the well-developed software LEfSe (Segata, Izard et al. 2011) to identify marker species, coincidentally, resulting in 40 markers (|LDA (Linear Discriminant Analysis) scores| >2) associated with T2D (Figure S1). 16 marker species are shared by the results of the NN model and LEfSe while the other 24 marker species are different (Table S5). However, with these marker species identified by LEfSe in D1 samples, the prediction accuracy of the NN model in D2+ samples became 74.7**%**. One possible reason may be that LEfSe ignores some important biomarkers with discriminant ability, the abundances of which show little changes across the case and control samples.

To characterize the variations of the 40 marker species identified by the NN model across the D1 samples, we compared the distribution of their relative abundances in case and control samples (Figure 2b). Based on the result obtained using the Kruskal-Wallis test, we can observe that only 16 of 40 marker species vary significantly between the case and control samples in their relative abundances (P-value < 0.05). Thus, we examined the impact of these 16 marker species on the classification performance and found that the average classification accuracy of the NN model (20 runs of 5-fold CV) for D1 samples using the 16 marker species become **93**.**6%**, which is down by almost 5% compared to the result of using the 40 marker species. Additionally, the classification accuracy of the model reduces to **73**.**6%** in testing on the D2+ dataset with these 16 marker species, suggesting that these species might not provide adequate discriminant information for the NN model in classifying the samples. Moreover, some of the marker species varying little across the case and control samples(e.g., *R. bromii* (ID: 40), *C. catus* (ID: 37), and *C. aerofaciens* (ID: 38)) have been detected associated with T2D in previous studies (Wu, Ma et al. 2010, Qin, Li et al. 2012, Murri, Leiva et al. 2013, Gurung, Li et al. 2020) (Table S8). This observation suggests that the marker species that do not vary significantly across the case and control samples likely play an important role in the gut microbial alteration related to T2D as well.

### The interaction network analysis identifies the key species in the alteration of the human gut microbiome

To further investigate the relationship among these marker species, we constructed the species-species interaction network based on the vector of the relative abundances of the marker species of D1. All possible interactions between species-species were ranked according to their reliability scores calculated by GENIE3, and we first selected the top 100 interactions for constructing the interaction network. As shown in Figure 2c, in total, 39 of the 40 marker species have at least one connection with others, 13 species connect with at least 5 other species (Figure 2d), and 4 species, including *Veillonella parvula, Streptococcus salivarius, Escherichia coli*, and *Escherichia_unclassified*, have more than 10 connections. Moreover, the relative abundances of 7 (green color) of the 13 species do not vary significantly across the case and control samples (Figure 2d). Among the 7 species, *E. coli* and *S. salivarius* have the highest number of connections, 35 and 25, respectively, suggesting the importance of these two species in the T2D-related alteration of the human gut microbiome. *E. coli* has been identified to be diabetes-enriched (Qin, Li et al. 2012) and *S. salivarius* can ferment glucose yielding lactic acid (Wescombe, Heng et al. 2009). Nearly 70% (n=9) of the 13 marker species (Figure 2d) is short_chain fatty acids-producing bacteria and previous studies have demonstrated that these types of bacteria played an important role in T2D (Qin, Li et al. 2012, Koh, De et al. 2016). Additionally, among the 13 species, except for *E*.*coli, B*.*vulgatus*, and *B. uniformis*, other species are extremely low abundant in the human gut samples. Thus, these findings from the interaction network analysis based on the selected marker species suggest that the dynamics of the T2D-related gut microbiota might be driven by few key players and those not varying significantly across the samples likely also play a critical role in the T2D-related alteration of the human gut microbiome by interacting with other species.

### Core functional gene modules related to T2D are identified by interaction network analysis of the marker genes

To identify the T2D-associated gut metagenomic markers on both genetic and functional levels, we also used our method on the functional gene profile of D1. Similarly, we ranked the genes using RF and then evaluated the average accuracies of the NN model with different ranked genes using 5-fold CV. As seen in Figure 3a, the highest value (**89**.**4%**) of the average accuracy was obtained when the top 90 genes were selected for classification. To this end, we took the 90 genes as the marker genes (Table S6).

**Figure 3.**
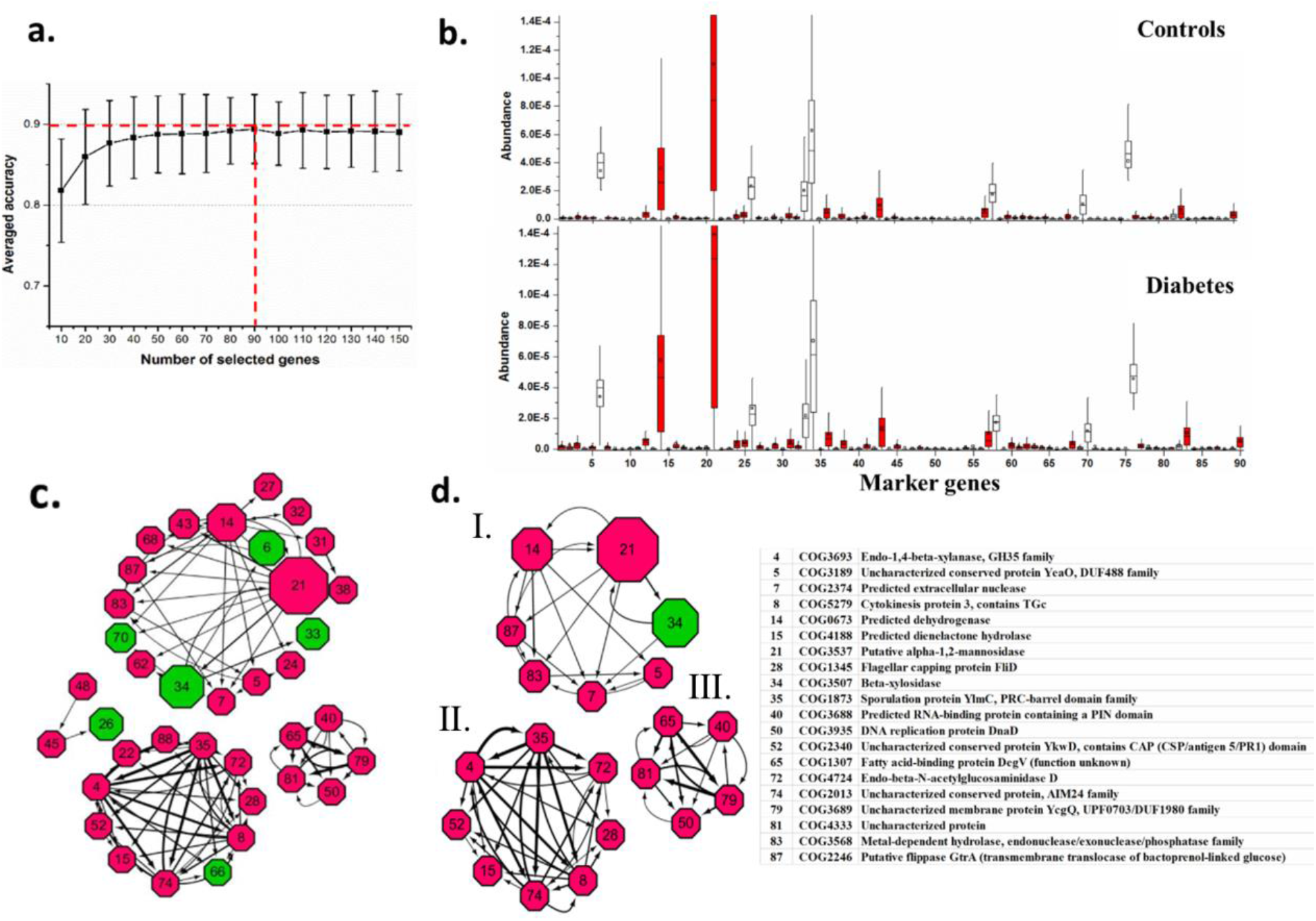
Marker gene identification and interaction network construction. **a**. Average accuracy of classification with the different number of the ranked genes based on Neural Network-based classifier by using 20 runs of 5-fold CV on the gene profile of D1. And the peak value could be obtained as the 90 species were selected, thus these genes were identified as the marker genes in this study. **b**. The distribution of the abundance of the marker genes that significantly varied across samples in the diabetes-related dysbiosis. A comparison between the cases (n = 183) and controls (n = 185) from the gene profile of D1. Using the Kruskal-Wallis test, the species with red color were identified that the abundances varied significantly (P-value < 0.05) across all samples. The order of the marker genes is the same as that in Table S6. **c**. The interaction network of marker genes determined by GENIE3, where the top 100 most reliable interactions between marker genes were selected. The width of the edges between nodes is proportional to the reliability of the linkage between the nodes. The size of each node is proportional to the average abundance of the marker genes. The network layout was calculated by the **Cytoscape** software using a circular layout. The number in each node is the ID of the marker species, which is consistent with Supplementary Table S2. The nodes with red color are the marker species that significantly were enriched in the diabetic samples or the control samples. **d**. Interactions between marker genes with a degree of more than 5.

We further tested the NN model using the 90-marker genes (trained on D1−dataset) in identifying the T2D samples of the D2+ dataset based on the gene profile, resulting in 88.6% classification accuracy. In the same way, we compared the relative abundances of these marker genes in case and control samples of the D1 dataset and observed that 78 out of the 90 genes vary significantly (P-value < 0.05) using Kruskal-Wallis test (Figure 3b). The average accuracy (20 run 5-fold CV) of the NN model using the 78 genes in classifying the samples of the D1 dataset reduces slightly from 89.4 to **88**.**3%**.

To characterize the interactions between the marker genes, we also constructed the gene-gene interaction network using GENIE3 (Figure 3c), resulting in 37 of 90 marker genes have at least one connection with other genes. As seen in Figure 3c, the interaction network of the marker genes is partitioned into four separate groups and the genes connect with others within each group. Few genes (n=20) have more than 5 connections with other genes (Figure 3d). We utilized BlastKOALA for annotation and KEGG mapping to characterize the functional category of the 90 marker genes in functional modules or pathways. As a result, we find 56.7% (n=51) of the marker genes can be annotated in the database COG (Figure S2). Only 16 of 90 genes can be mapped to the functions in the KEGG database. According to the functional annotations of these marker genes in the KEGG database, 4 of 18 genes in group I (ID:34(COG3507), 70(COG2160), 87(COG2246), 68(COG1071)), 2 of 3 genes in group II (ID: 48(COG 1211), 26(COG0493)), 1of 11 gene in group III (ID:66(COG2971) were mapped to the pathways of carbohydrate metabolism (Table S7), suggesting these genes in the groups are likely involved in or associated with the carbohydrate metabolism. One of 11 genes in group III (ID: 28(COG 1345) was mapped to flagellar assembly. The marker genes connect with others in each group but do not interact with those in other groups, implying the genes in different groups function differently in carbohydrate metabolism. The interaction of these genes confirms the link between the T2D-related alterations of the human gut microbiome and carbohydrate metabolism(Hammes, Du et al. 2003, Koh, De et al. 2016, Zhao, Zhang et al. 2018).

### The NN-based regression analysis determines the most significant covariate factor associated with the T2D-related microbiome alteration

FBG (fasting blood glucose), age, BMI (body mass index), and the weight of patients all potentially are causative factors of T2D. To determine the most significant factors associated with the human gut microbiome represented by the T2D-related microbial markers, we did the regression analysis of D1 with the 40 marker species and the 90 marker genes, respectively. Based on the 5-fold CV, for each factor, the predicted values of the samples were calculated by the NN model. Then, these values and the corresponding real values were plotted (Figure 4). These figures demonstrate that for each factor, the microbiome values predicted based on the markers are linearly correlated with the corresponding real values across the samples (Figure 4 a, b, c, d, and Figure S3; P<<0.05), suggesting that the human gut microbiome covaries with these factors. In particular, the microbiome correlation with FBG (P< e-50) is more significant than with other factors (age, 1e-43; BMI,1e-30; Weight, 1e-32), indicating that FBG is likely the leading factor in driving the T2D-related alteration of the human gut microbiome in the development of T2D. This result is in line with that of the analysis on the gene-level (Figure S3, R=0.686, P=2.00e-52). Although several studies (Suez, Korem et al. 2014, Korem, Zeevi et al. 2017) have reported the relationship between the gut microbiome and FBG, our result reveals the strong correlation between FBG and the T2D-related alteration of the human gut microbiome using the non-linear regression rather than classification.

**Figure 4.**
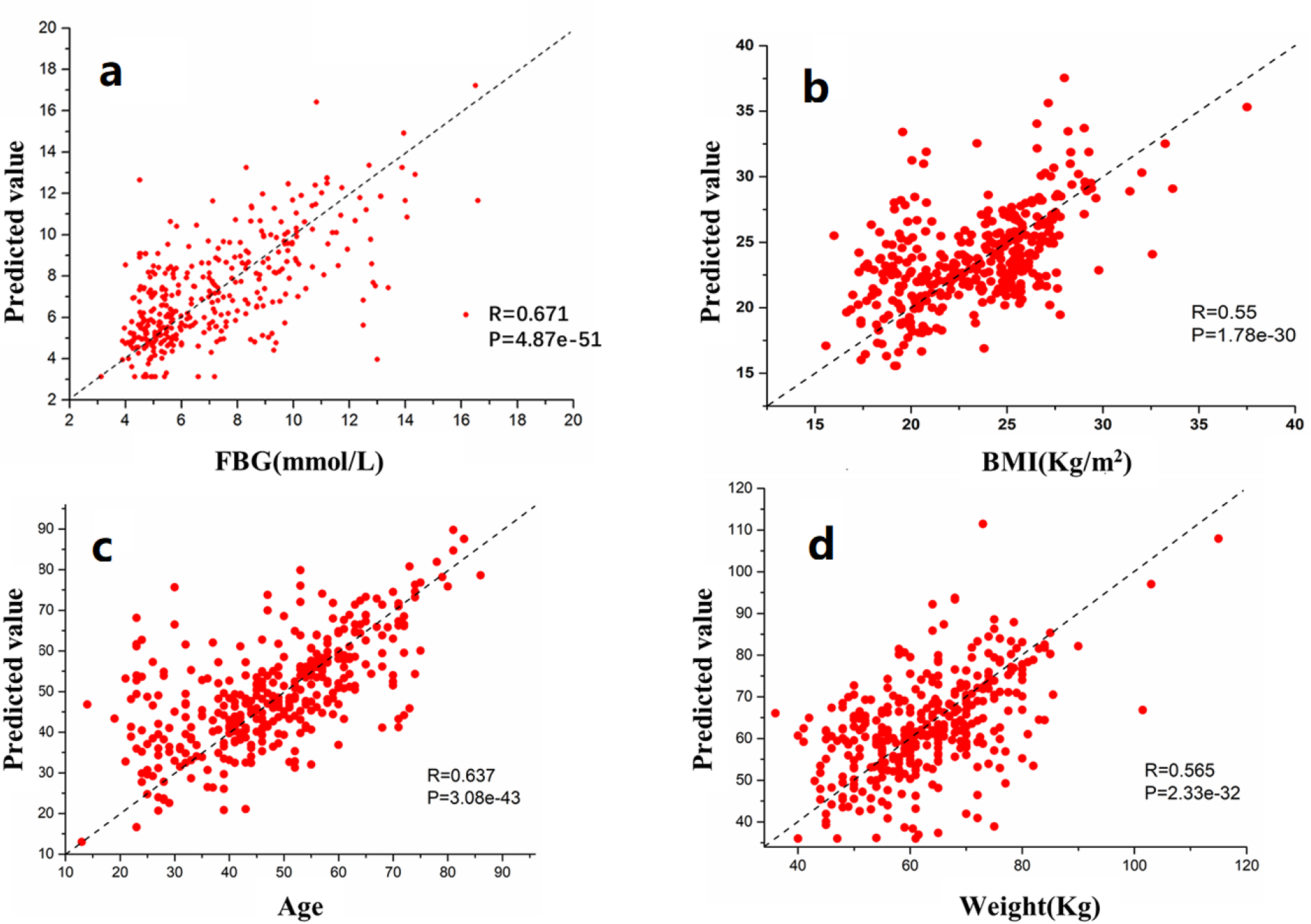
Neural Network-based predicting values and the actual values of T2D-related factors (i.e., FBG, age, weight, BMI) based on the species level profile of D1 with our marker species. Each T2D-related factor values were fit by the corresponding NN based regression model using 5-fold CV, and the R-value (i.e., Pearson’s linear correlation coefficient), as well as the P-value between the real values and the predicted values, were obtained by statistical calculation.

To explore how these marker species covary with the dynamic change of FBG in the development of T2D, we plotted the heat map of the mean relative abundances of the marker species on four intervals (i.e., Q1: < 5.02; Q2: 5.02∼6.21; Q3: 6.21∼8.8; Q4: > 8.8) of FBG (Figure 5). It can be observed that the relative abundances of these marker species vary greatly at different intervals (Figure 5). Among the 16 marker species with a significant difference in relative abundance between case and control samples (P-value < 0.05) based on the Kruskal-Wallis test, 6 have higher relative abundances in healthy control samples, 10 in T2D samples. Those 6 marker species (Figure 5, black) are higher abundant in both Q1 and Q2 than in Q3 and Q4 samples. Among those 10 marker species, 6 are higher abundant in both Q3 and Q4, 1(*B*.*dorei*) in Q4, 1 (*E*.*lenta*) in Q3, and 2 (*Veillonella. unclassified* and *S. anginosus*) in both Q1 and Q3. These marker species are identified as T2D-related species easily according to their relative abundances in T2D (high FBG) and healthy samples (low FBG) using conventional statistic-based methods. Other 24 marker species that do not have significant differences in relative abundance between the case and control samples exhibit diverse patterns. Intuitively, some of them (n=3, *D*.*longicatena, P*.*copri*, and *S*.*salivarius*,) reach to the highest abundance in Q1, 2 (*B*.*ovatus, Escherichia*.*unclassified*) in Q2, 5 (*V*.*parvula, B*.*xylanisolvens, E*.*eligens, R. bromii, B*.*longum*) in Q3, 3 (*A*.*shahii, E*.*coli*, and *E*.*ventriosurn*) in both Q2 and Q3, 2 (*B*.*thetaiotaomicron, E*.*hallii*) in both Q2 and Q4. Thus, these 24 marker species vary in different patterns associated with the dynamic changes of FBG. This may explain the reason that the two important T2D-related marker species (*E*.*coli and S*.*salivarius*) detected by the interaction network analysis vary little across the case and control samples. In conclusion, our analysis suggests that these marker species are likely to play a different role in different stages of the T2D development.

**Figure 5.**
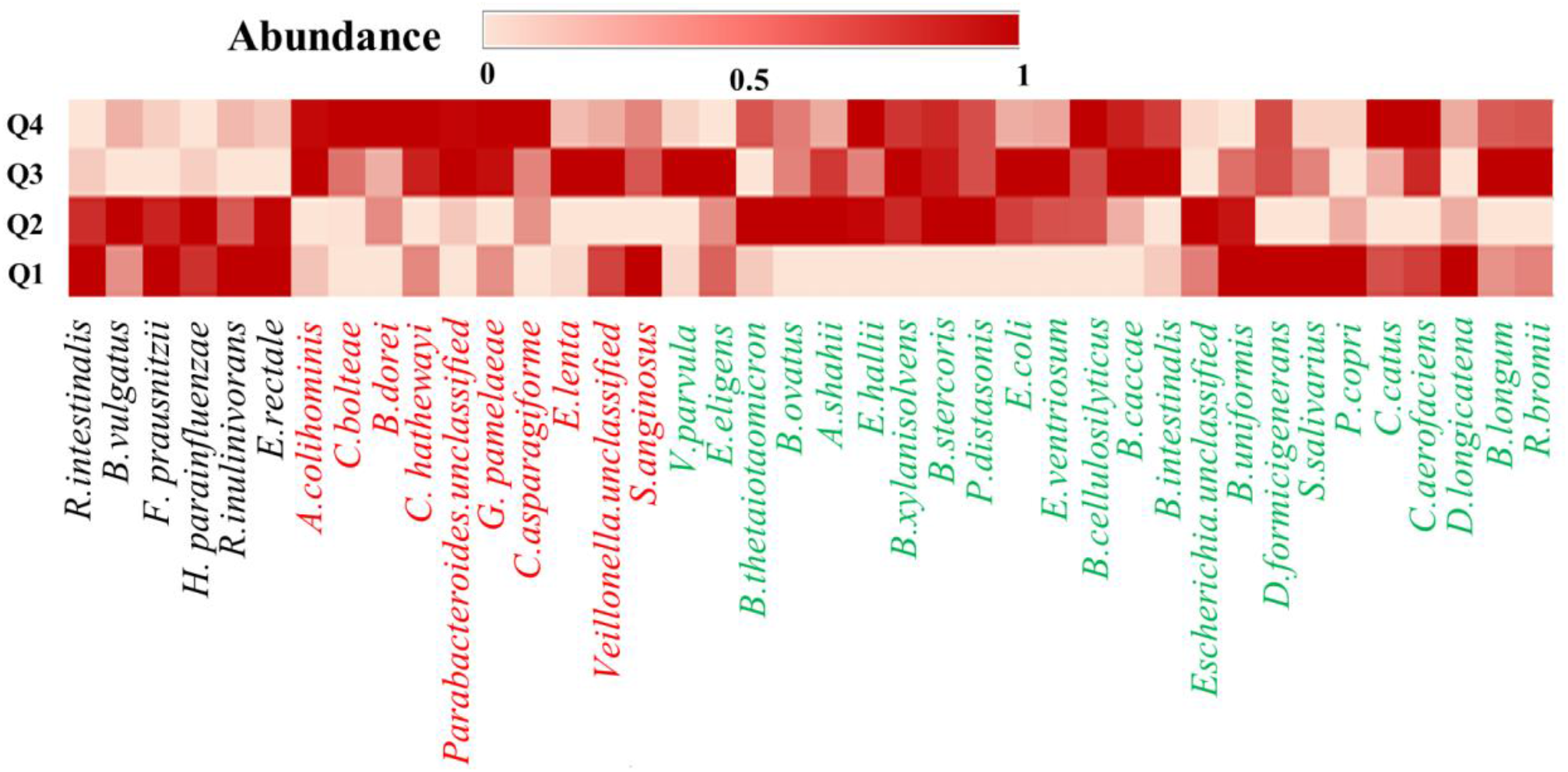
Heat map of mean relative abundances of the 40 selected marker species on the four dynamic intervals of fasting blood glucose. The dynamic intervals were determined by the quantile statistics (Q1:< 5.02; Q2: 5.02∼6.21; Q3: 6.21∼8.8; Q4: > 8.8). In each dynamic interval, the abundances of marker species for the corresponding samples were averaged. Then, the average abundances of each marker species for the four dynamic intervals were normalized (mapping to [0, 1]). The species names were colored based on the significance analysis across samples using the Kruskal-Wallis test (black: abundances significantly increase in healthy samples; blue: abundances significantly increase in T2D samples; green: abundances have not significant differences between T2D and healthy samples).

## DISCUSSION

To provide different perspectives for understanding the roles of the gut microbiota in the development of T2D, here we present the NN-based framework to identify microbial markers that can be used as representatives of the gut microbiota for prediction of T2D-related samples with high performance. Many markers not detectable by conventional statistic methods vary little across the case and control samples. Nevertheless, interaction network analysis and regression analysis indicate that these markers likely play a crucial role in the T2D-related alteration of the human gut microbiome by interacting with other markers. This framework firstly uncovers the T2D-related dynamics and interaction of the microbial markers in the human gut microbiome, strongly suggesting the cumulative effect of the markers rather than individuals is likely the driver of the gut microbiome alteration.

The deep learning approach usually requires massive samples for training and feature extraction. Although numerous T2D-related human gut microbiome studies have been completed (Gurung, Li et al. 2020), we only recruited the datasets generated by whole genomic sequencing and thus eligible for this study (Qin, Li et al. 2012, Zhao, Zhang et al. 2018). The main limitation of our study is that only a small number of samples (N=368, D1) were recruited for training the NN model. Even so, by determining the suitable numbers of layers and nodes, our model obtained higher performance in classifying the T2D-related samples in the validation experiment than other approaches. We also used our model (trained on D1, Chinese samples) using the identified markers for diagnosing T2D in the dataset D3 of European women (Karlsson, Tremaroli et al. 2013), however, resulting in the low accuracy of 52.1% and 57.4% on species- and gene-level, respectively (data not shown). This reflects the great variation of the human gut microbiome of the subjects living in geographically and culturally distinct settings (Yan He 2018). Thus, here we only show the results based on the datasets of D1 and D2 in the study.

The classification results demonstrate that our NN model outperforms much better than other conventional machine-learning methods including SVM-linear, SVM-RBF, RF, and KNN (Figure 1b), showing its high capability to novel patterns in complex microbiome datasets (Pasolli, Truong et al. 2016). Our analysis demonstrates that the NN model has higher performance in classifying the T2D-related samples using our selected markers than the markers identified by LEfSe. The interactions among these marker species may carry more information than the individual species. In this regard, the marker species that do not vary significantly in relative abundance across the case and control samples likely play a special role in the gut microbial alteration related to T2D as well. The software LEfSe does not take the interaction between the species into account, while our method takes non-trivial relationships in the data into account and therefore will produce better answers.

Regression analyses show that the human gut microbiome represented by the selected microbial markers has a strongest correlation with FBG in terms of the R and P values (R=0.678, P=4.87e-51, Figure 4). This finding indicates that the gut microbial composition of patients likely keeps altering with the increase in the value of FBG in the development of T2D. We stratified the samples of D1 into four intervals according to their FBG values (i.e., Q1: < 5.02; Q2: 5.02∼6.21; Q3: 6.21∼8.8; Q4: > 8.8) and plotted the heat map of mean relative abundances of the marker species of the samples within each interval. We can observe that the 24 marker species that do not vary significantly across the case and control samples have different patterns in relative abundances across the 4 intervals of FBG values, showing that they are as important in driving the T2D-related gut microbiome alteration as those with significant differences between case and control samples identified by statistic-based methods. These results provide a different perspective for understanding the relationship between diabetes and the human gut microbiome and thus need further investigations.

In summary, to our knowledge, for the first time, this study develops a framework combining NN and RF to re-analyze the human microbiome data to identify the T2D-related microbial markers. Our results demonstrate that the interaction of these markers carries more information than the individuals in predicting the disease state of the samples, suggesting that the NN model can help us take non-trivial relationships in the complex microbiome data into account. This study paves a way to use the NN algorithm in microbiome study and provides potential opportunities to deeply understand the microbial roles in the development of human diseases and evaluate individuals at the risk of relevant diseases.

## Abbreviations

T2D: type II diabetes
NN: neural network
RF: random forest
FBG: fasting blood glucose
CV: cross-validation
ROC: receiver operating characteristic
AUC: area under the curve
BMI: body mass index

## Declarations

### Ethics approval and consent to participate

N/A

### Consent for publication

N/A

### Availability of data and materials

The data (i.e., the profiles of D1 and D2) and the demo of the NN model is available in the GitHub repository (https://github.com/gsgowell/microbial_markers_identification).

### Authors’ contributions

Y.M and S.G designed the research. S.G., Y.C., H.Z., and S.J. performed analysis. S.G., and Y.M., drafted the paper. All authors contributed to the interpretation of the results and the text.

## Acknowledgments

N/A

## Supplementary Materials

Table S1 Parameters of the species-based classifier

Table S2 Parameters of the gene-based classifier

Table S3 The prediction performance of the classifier with different number of hidden layers based on the species level profile of D1

Table S4 The list of diabetes-associated gut microbiota species that selected by our method

Table S5 The list of diabetes-associated gut microbiota species that were selected simultaneously by our method and LEfSe.

Table S6 The list of diabetes-associated gut microbiota genes that selected by our method

Table S7 The functional prediction of the marker genes in KEGG

Table S9 The list of the marker species that could be assigned to previously known diabetes-related species or genus

Figure S1 The marker species identified by LEfSe from D1

Figure S2 The functional category of the marker genes on the module or pathway level by using BlastKOALA.

